# Crystal structure of a MarR family transcriptional regulator protein

**DOI:** 10.1101/2024.06.07.597875

**Authors:** Arpita Goswami, Kannika Byadarahalli Ravindranath, Madan Kumar Shankar

## Abstract

The multiple antibiotic resistance regulator (MarR) family of transcription factor proteins form a large group of multitasking bio-molecules in pathogenic *Escherichia coli* (*E. coli*). HosA is one of these MarR transcription factors reported in dozens of pathogenic *E. coli* with highly conserved sequence profiles. The HosA from the enteropathogenic *E. coli* O127:H6 (strain E2348/69), a predominantly monomeric protein, was overexpressed in *E. coli* and purified. The HosA protein crystals were obtained in microbatch under oil method at 4° C. The X-rays of the diffracted spots were extended to 2.21 Å resolution. The crystal belongs to the space group *P*4_3_22, with unit-cell parameters *a* = 67.16 Å, *b* = 67.16 Å, *c* = 95.66 Å and α = β = γ = 90°. In the asymmetric unit, monomeric HosA protein was crystallized and confirmed with the Matthew coefficient analysis (3.48 Å^3^ Da^-1^). The monomeric structure is compared with previously solved structures of other homologous transcription factors. This confirmed the winged loop at the DNA binding region of the HosA protein.

## Introduction

In nature, protein molecules often evolve through gene duplication events in responses to environment stimuli (Lynch & Conery, 2000). The multiple antibiotic resistance regulator (MarR) family of transcription factor regulator proteins are such gene duplication products involved in antibiotic resistance (Cohen, 1993), virulence (Ellison & Miller, 2006), central metabolism (Grove, 2017; Huang *et. al*., 2015), metal transport (Deochand & Grove, 2017), oxidative stress (Fuangthong & Helmann, 2002; Deochand & Grove, 2017), degradation of aromatic compounds (Deochand & Grove, 2017) among others. Most of these proteins act as inhibitors for self transcription as well as target operons. However, sometimes they are also involved in activations of target genes by attenuating methylation H-NS mediated inhibition, as seen in SlyA (Wyborn, 2004). HosA (Homologue of SlyA) is another SlyA-like MarR transcription factor reported in dozens of pathogenic *Escherichia coli* (*E. coli*) with highly conserved sequence profiles (Ferrandiz et al, 2005). The auto-inhibition of HosA by binding to DNA is reported for uropathogenic *E. coli* UMN026 (Roy & Ranjan, 2016). In the same study, HosA also repressed non oxidative hydroxy acrylic acid decarboxylase (HAD) operon. The enzymes produced by this operon primarily act on 4-hydroxy benzoic acid (4-HBA) or paraben (Roy & Ranjan, 2016). Therefore, presence of paraben in the environment caused the reversal of this inhibition through direct binding to HosA. Similar allosteric regulation mediated by organic molecules binding to conserved sites has also been reported for other MarR regulators (Deochand & Grove, 2017; Perera & Grove, 2010). In addition to its involvement in paraben metabolism, this protein also functions as an activator for temperature dependent flagellar motility by directly binding to flagellin DNA promoter in enteropathogenic *Escherichia coli* (Ferrandiz et al, 2005). All these diverse functions of HosA protein may be easier to explain in presence of structural information which is absent in literature. Therefore in this study, the HosA protein from enteropathogenic *E. coli* O127:H6 (strain E2348/69) is purified, crystallized for X-ray diffraction and ensuing structural studies were carried out. This work may provide a platform for unravelling the allosteric regulation of HosA and its interactions with DNA and organic molecules.

## 2. Materials and methods

### 2.1. Macromolecule production

The *Escherichia coli* O127:H6 (strain E2348/69) HosA DNA sequence retrieved from NCBI (Reference genome: NZ_LT827011.1) was codon optimized for *Escherichia coli*. The sequence was further cloned in pET21a+ (Novagen) between NdeI and XhoI restriction sites from Genscript, USA. The synthesized gene had a 6X-His tag from the vector at C-terminus before the stop codon. Further, the cloned plasmid was transformed into BL21(DE3) cells (Novagen). Primary culture in LB (Luria Bertani) broth (HiMedia) from the transformed colony was grown overnight at 37°C and 200 rpm. 5 ml of secondary LB culture from the same (1 % inoculum) was grown at 37°C until OD reached 0.6. 1 mM IPTG was added thereafter and further the culture was grown for 4 hours at 37°C. In parallel, 5 ml uninduced culture was maintained using the same conditions but without IPTG. The overgrown cultures were pelleted, resuspended in a lysis buffer (50 mM Tris-Ph 8, 150 mM NaCl, 100 μM PMSF) and lysed by sonication. Post sonication lysates were centrifuged at 12,000 rpm at 4°C and 12% SDS-PAGE gel was run to observe solubility profiles of the protein after expression. The overexpression of the culture was done in 2 liter LB broth using optimized expression conditions as described above. The supernatant from lysed & centrifuged cell pellets was loaded in the Ni-NTA column (HisTrap FF, Cytiva). The washing and purification was done as per manufacturer’s protocol. The eluted protein in one fraction was >99% pure. This fraction was concentrated to 500 *µ*l and used for crystallization. The final concentration of the protein as estimated from the BCA test (Sigma Aldrich) was 8.4 mg/ml.

**Table 1.**
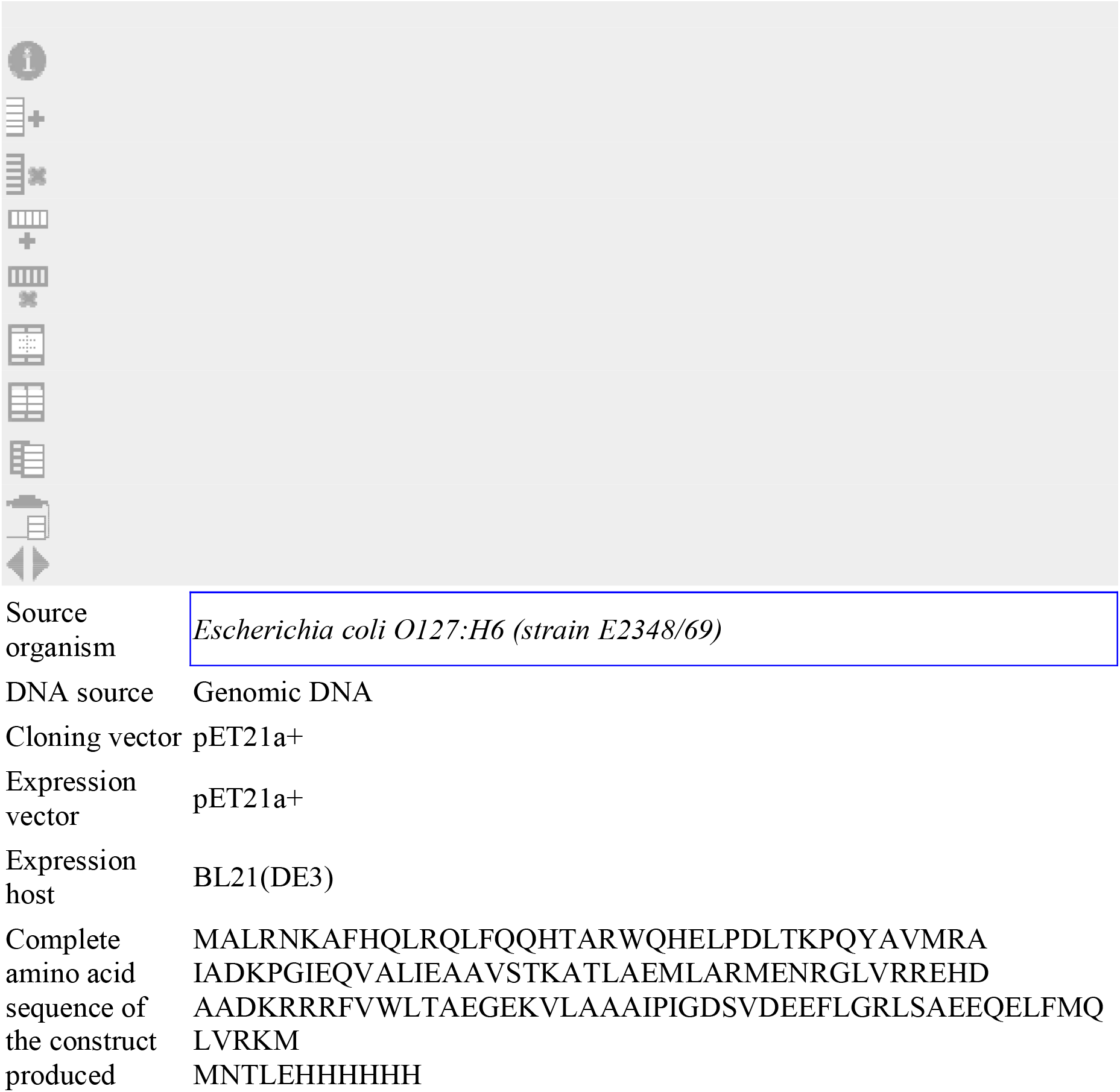
Macromolecule production information. In the primers, indicate any restriction sites, cleavage sites or introduction of additional residues, *e*.*g*. His6-tag, as well as modifications, *e*.*g*. Se-Met instead of Met.

### 2.2. Crystallization

The apo-protein was crystallized in microbatch under oil method using 96 well vapor batch plates (Douglas) as 1:1 ratio (protein:precipitant). The usual screens from Hampton, Rigaku and Jena were used. Best quality crystals were obtained in 4 °C and low pH conditions with ammonium acetate and PEG3350. The conditions were further optimized by grid setup. Final diffraction quality crystals were grown in 160 mM ammonium acetate, 100 mM Bis-Tris (pH 5.6), 27% PEG3350 (w/v). The concentration of protein was 6 mg/ml for all crystallization experiments.

**Table 2.** Crystallization.

|  |  |
| --- | --- |
| Method | Microbatch under oil |
| Plate type | Douglas vapour diffusion plate |
| Temperature (K) | 277.15 |
| Protein concentration | 6 mg/ml |
| Buffer composition of protein solution | 10 mM Tris HCl (pH 8) |
| Composition of reservoir solution | 160 mM ammonium acetate, 100 mM Tris HCl (pH 5.6), 27 %, PEG |
| Volume and ratio of drop | 1:1 |
| Volume of reservoir | 1 ul |

### 2.3 Data collection and processing

The crystals were scooped in cryoloop (Hampton) with 25% ethylene glycol as cryoprotectant added to the respective mother liquors. After flash freezing in liquid nitrogen, the crystals were subjected to diffraction in Rigaku Micromax-007 HF diffractometer with MAR 345 image plate detector and rotating anode X-ray generator set at 50 kV, 100 mA voltage. Preliminary space group analysis in iMosflm (ver. 7.4.0) (Winn et al., 2011) from the first few frames collected showed the space group as P4 for apo-protein crystals. Accordingly, the data collection tasks were set up to get >99% completeness in ∼ 5 hours. Indexing at first pointed to the P4_2_11 group by Pointless (ver. 1.12.12; iMosflm) followed by scaling in Aimless & Ctruncate (ver. 0.7.7 & 1.17.29; iMosflm) (Winn et al., 2011) without any crystal defects (Twinning, Pseudo-translation etc.). Since, the presence of screw axis led systematic absences are known in this group. Further correction was obtained while doing molecular replacement in Phaser (ver. 2.8.3, CCP4i-7.1.018) (Winn et al.,2011) by selecting analysis of all possible groups in P4. This conveniently confirmed the space group as P4_3_22. The Pointless indexed reflections were then reindexed (Reindex, CCP4i, ver. 7.1.018) to P4_3_22, scaled (Scala, CCP4i, ver. 3.3.22) and this data was used in refinement later by phenix.refine (Phenix, ver. 1.19.2_4158) to prevent model bias. The Table 3 shows values from this reprocessed data.

**Table 3.** Data collection and processing. Please use the footnotes section beneath this table to address any issues highlighted in <u>**bold-underlined**</u> text if applicable Values for the outer shell are given in parentheses.

|  |  |
| --- | --- |
| Diffraction source | RIGAKU MICROMAX-01 |
| Wavelength (Å) | 1.54179 |
| Temperature (K) | 100 |
| Detector | MAR 345 image plate |
| Crystal-detector distance (mm) | 200 |
| Rotation range per image (°) | 1 |
| Total rotation range (°) | 89 |
| Exposure time per image (s) | 120 |
| Space group | $P4_322$ |
| $a, b, c$ (Å) | 67.16, 67.16, 95.66 |
| $\alpha, \beta, \gamma$ (°) | 90, 90, 90 |
| Mosaicity (°) | 1.33 |
| Resolution range (Å) | 54.97-2.21 (2.33-2.21) |
| Total No. of reflections | 76419 (9702) |
| No. of unique reflections | 11402 (1524) |
| Completeness (%) | 99 (93.3) |
| Redundancy | 6.7 (6.4) |
| $\langle I/\sigma(I) \rangle$ | 9.3 (2.3) |
| $R_{\text{r.i.m.}}$ | 0.152 (0.827) |
| Overall $B$ factor from Wilson plot (Å<sup>2</sup>) | **<u>15.204</u>** |

### 2.4. Structure solution and refinement

The template in Phaser was the C chain from the hexameric model of 6PCP (*Bordetella bronchiseptica* BpsR with 28% identity to HosA, RCSB PDB) (Booth et al., 2019). The single solution obtained from Phaser was subjected to model building in Autobuild (Phenix, ver. 1.19.2_4158) (Liebschner et al., 2019) using multiple non-crystallographic symmetry (ncs) options (= 1, 2, 3). A well-built model was obtained only in ncs=3 after a single cycle of model building. There was also high water content in the crystal (64.67% as determined by Mathew coefficient calculation as monomer in asymmetric unit) (Matthews, 1968). . The corresponding value of Mathew coefficient was 3.48. Final model for refinement was obtained after multiple cycles of model building from this initial model with ncs =1. As mentioned before, reindexed and scaled reflections were used for refinement of coordinates in phenix.refine (Phenix, ver.1.19.2_4158). The accuracy of the reindexed intensities were further verified by cross-checking in Xtriage (Phenix, ver. 1.19.2_4158) which otherwise showed anomalies in original indexed data. The refinement was carried out until there were no structural defects and good refinement values (R_work_/R_free_ = 0.21/0.26). The final refinement results are detailed in Table 4.

**Table 4.** Structure refinement. Please check <u>**bold-underlined**</u> values (these may have been derived because they are not explicitly defined in the CIF) Values for the outer shell are given in parentheses.

|  |  |
| --- | --- |
| Resolution range (Å) | 47.49–2.25 (2.48–2.25) |
| Completeness (%) | 100.0 |
| $\sigma$ cutoff | $F > 1.35\sigma(F)$ |
| No. of reflections, working set | 10925 (2529) |
| No. of reflections, test set | 546 (130) |
| Final $R_{\text{cryst}}$ | 0.212 (0.2477) |
| Final $R_{\text{free}}$ | 0.258 (0.3235) |
| No. of non-H atoms | 1128 |
| Protein | 1023 |
| Ligand | **<u>4</u>** |
| Water | 101 |
| R.m.s. deviations |  |
| Bonds (Å) | 0.002 |
| Angles (°) | 0.482 |
| Average $B$ factors (Å<sup>2</sup>) | |
| Protein | **<u>31.05</u>** |
| Ramachandran plot | 97.66 |
| Most favoured (%) | 2.34 |
| Allowed (%) | NIL |

## 3. Results and discussion

### 3.1 Protein expression and crystallization

The expression test was carried out from primary LB culture inoculated with a single transformed BL21(DE3) colony. 1% inoculum from overgrown primary culture was transferred to 5 ml LB media. This was grown at 37°C till mid log phase (OD 0.6). After that, 1 mM IPTG was added and induced culture was grown for further 4 hours. Similarly, uninduced secondary culture was maintained. The pellets obtained from both uninduced and induced secondary cultures were lysed, centrifuged and SDS-PAGE gel was run (Fig. 1a(i)). The molecular weight of the HosA protein band observed from all SDS PAGE runs was slightly less than 17 kD. This was close to the theoretical molecular weight (∼16.6 kD) calculated from the Peptide 2.0 server. It was also observed that HosA protein gets fractionated in supernatant after centrifugation of lysed culture.

**Figure 1.**
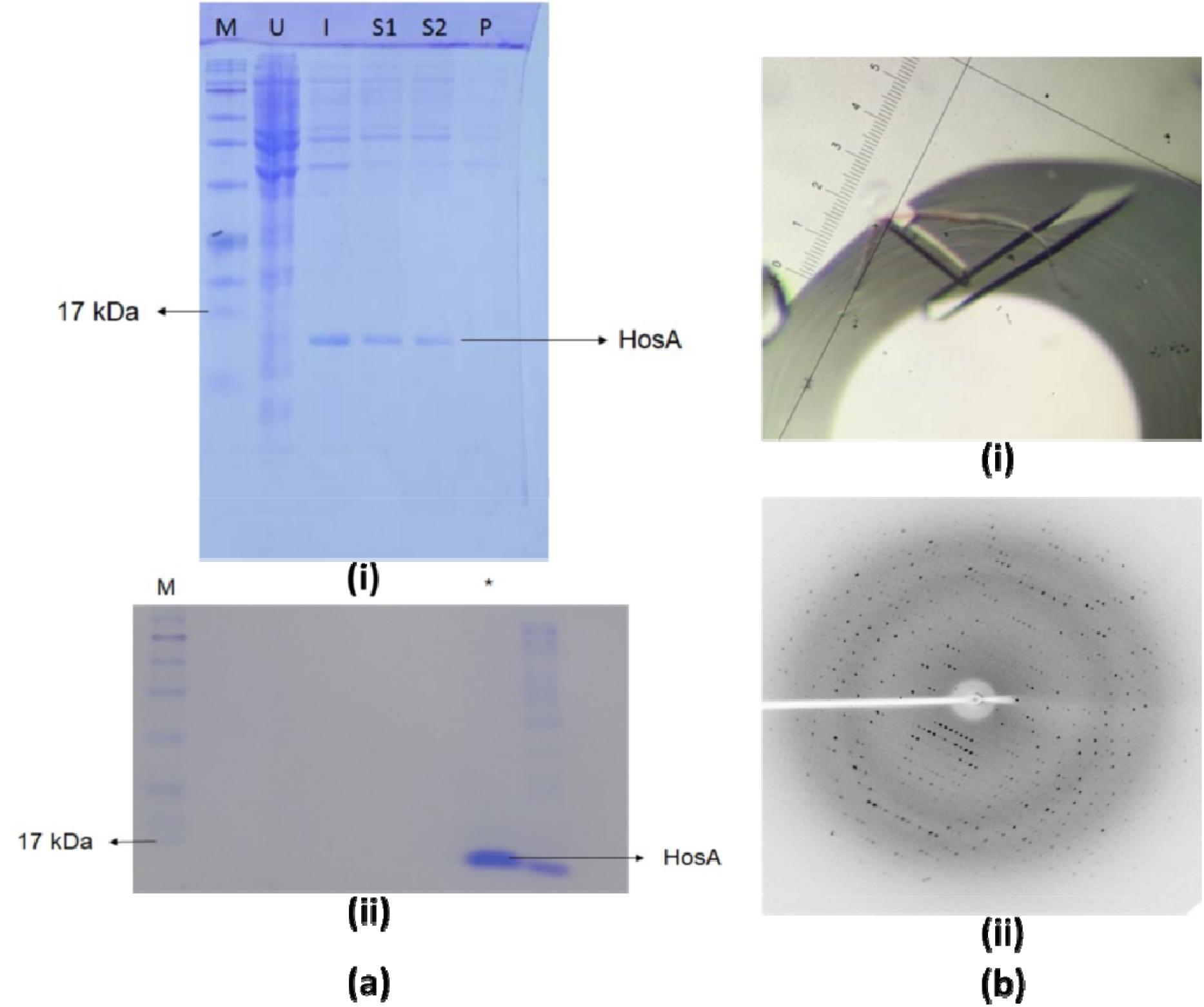
(a) SDS PAGE gels: (i) expression test from 5 ml cultures. U = Uninduced whole cell lysate, I = Induced whole cell lysate, S1 & S2 = Supernatants from induced whole cell lysate after centrifugation, P = Pellet from induced whole cell lysate after centrifugation and M = Jena BlueEye Prestained Protein Marker (10-245 kDa) (ii) Ni-NTA purification from 2 liters overexpressed culture. The fraction showing >99% purity is marked by *. M = Jena BlueEye Prestained Protein Marker (10-245 kDa). The protein band comes below 17 kDa in both gels. (b) Crystal for data collection and (ii) diffraction image. The bar in b (i) = ∼1 mm corresponds to scale.

The presence of protein in the insoluble fraction was negligible. This was also noted earlier for HosA expressed from uropathogenic *Escherichia coli* UMN026 (Roy & Ranjan, 2016). Nevertheless, this paved the way for further overexpression of the protein and other studies. As expected, the protein obtained from overexpressed culture also behaved similarly with almost no traces of protein in the insoluble fraction. Also, after purification with Ni-NTA chromatography the protein came as very pure with >99% purity in one eluted fraction (Fig. 1a(ii)).

The crystallization trials of HosA gave rod shaped crystals at 4°C in microbatch under oil method using 96 well Douglas vapor batch plates as 1:1 ratio (protein: precipitant, protein concentration 6 mg/ml)). The diffraction quality crystals (Fig. 1b (i)) after grid set up appeared in ammonium acetate (160 mM), Bis-Tris (100 mM, pH 5.6), 27% PEG3350 (27% w/v). The overall size of the biggest crystal was ∼ 6 mm X 0.5 mm (Length X Width) (Fig. 1b (i)). This crystal diffracted up to 2.21 Å resolution [Fig. 1b (ii)] in Rigaku Micromax-007 HF diffractometer and data was collected.

### 3.2 Monomeric structure and structure similarity

The structure of the HosA of *E. coli* (Fig. 2) was determined to 2.25 Å resolution and the asymmetric unit of the crystal structure has monomer as supported with the Matthews coefficient value of 3.48 Å^3^ Da^-1^ (Matthews, 1968). The overall monomeric structure has 6 □-helices (*α*1, *α*2, *α*3, *α*4, *α*5 and *α*6) and winged *β*-sheet formed by 3 antiparallel strands (tiny *β*1, longer *β*2 and *β*3) as observed across MarR family transcription factors. The sequence of strands are *α*1-*α*2-*β*1-*α*3-*α*4-*β*2-*β*3-*α*5-*α*6. In contrary, PISA analysis showed the molecule with 2 subunits in two asymmetric units of the unit cell to correlate with each other by a 2-fold crystallographic rotation axis. The total solvent accessible surface area for each monomer was 9099.5 Å^2^. This was reduced by 2223.3Å^2^ (buried surface area) upon homodimer formation. So total buried area for the dimer was 4446.6 Å^2^, sufficiently large to support assembly formation. The total solvent accessible surface area in the dimer was 13752.4 Å^2^. The solvation free energy gain upon formation of the assembly (ΔG^int^) was -35.6 kcal/mole indicating energy favorable association. While the free energy of assembly dissociation (ΔG^diss^) and the rigid-body entropy change at dissociation (TΔS^diss^) was 33.6 kcal/mol and 12.3 kcal/mol respectively indicating thermodynamic stability. Therefore, a stable crystallographic dimer upon close association of molecules is possible.

**Figure 2.**
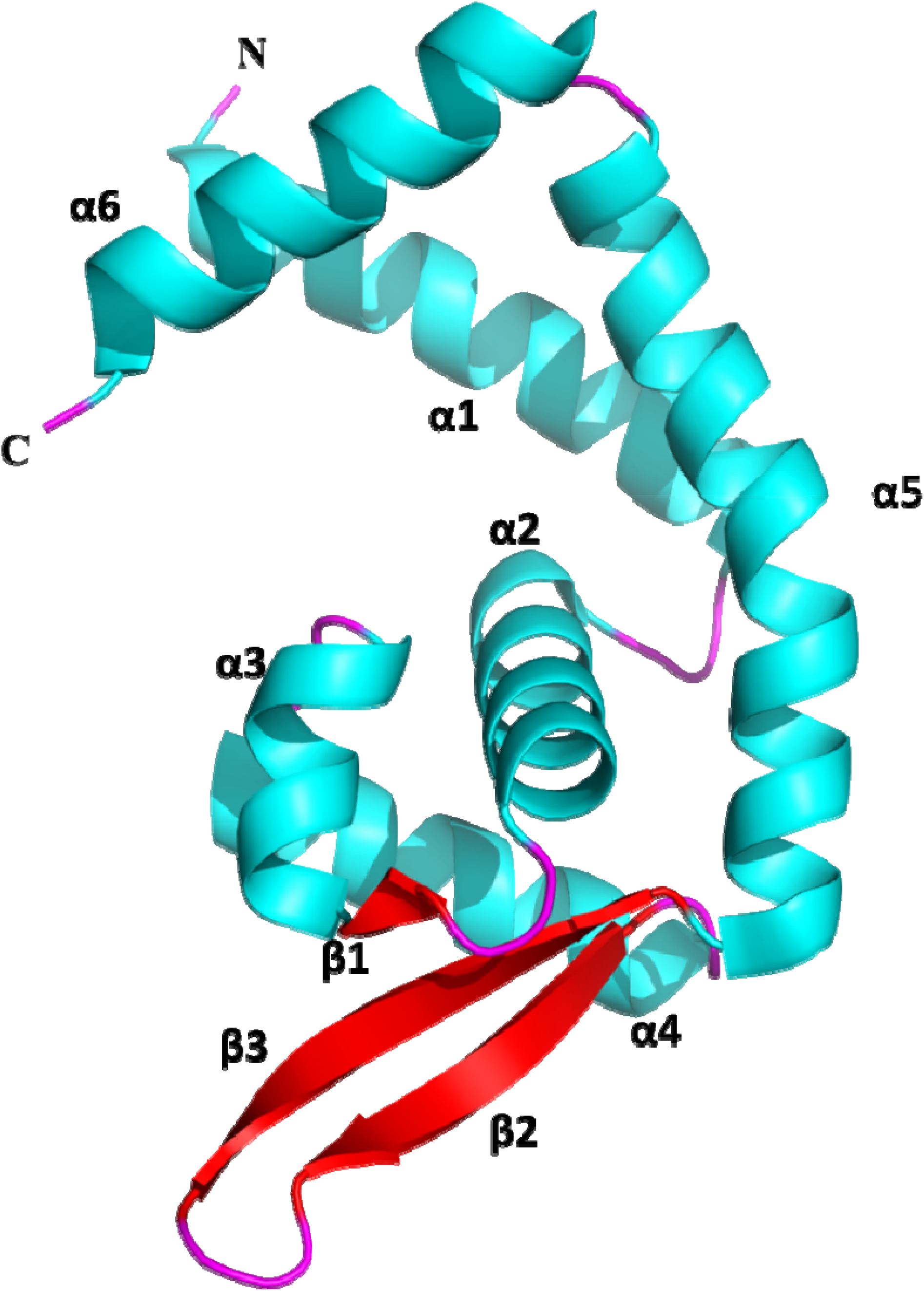
The overall monomeric structure (cartoon representation; cyan helices, red beta strands and pink loops) of HosA (8PQ4). The N-terminus and C-terminus are marked.

A DALI server search of the HosA monomer (PDB code 8PQB) against the Protein dataBank reveals a strong conservation of secondary structural elements with other transcriptional regulators (Fig. 3a and Fig.3b) based on PDB90 benchmark. The top closest homologs include different transcription factors from *Pseudomonas Aeruginosa* (PDB code 2NNN, Z-score 17.1, RMSD of 1.9 Å over 129 aligned C*α* atoms and sequence identity of 26 %),*Bordetella Bronchiseptica* (PDB code 6PCP, Z-score 17.1, RMSD of 1.6 Å over 130 aligned C*α* atoms and sequence identity of 29 %), *Streptomyces coelicolor* (PDB code 4FHT, Z-score 16.6, RMSD of 1.9 Å over 130 aligned C*α* atoms and sequence identity of 32 %), *Rhodopseudomonas palustris* (PDB code 6C2S, Z-score 16.1, RMSD of 2.0 Å over 130 aligned C*α* atoms and sequence identity of 19 %), *Acinetobacter Baumannii* (PDB code 7EL3, Z-score 16.1, RMSD of 2.0 Å over 130 aligned C*α* atoms and sequence identity of 18 %), *Pseudomonas putida* (PDB code 7KUA, Z-score 16.0, RMSD of 2.1 Å over 130 aligned C*α* atoms and sequence identity of 16 %) and *Escherichia coli K-12* (PDB code 5H3R, Z-score 15.9, RMSD of 2.1 Å over 130 aligned C*α* atoms and sequence identity of 24 %). Overall, the aligned loop regions of these superposed structures (Fig. 3b) clearly suggests the comparative uniqueness of conformation of the winged loop of HosA and this region is mainly associated with the DNA binding.

**Figure 3.**
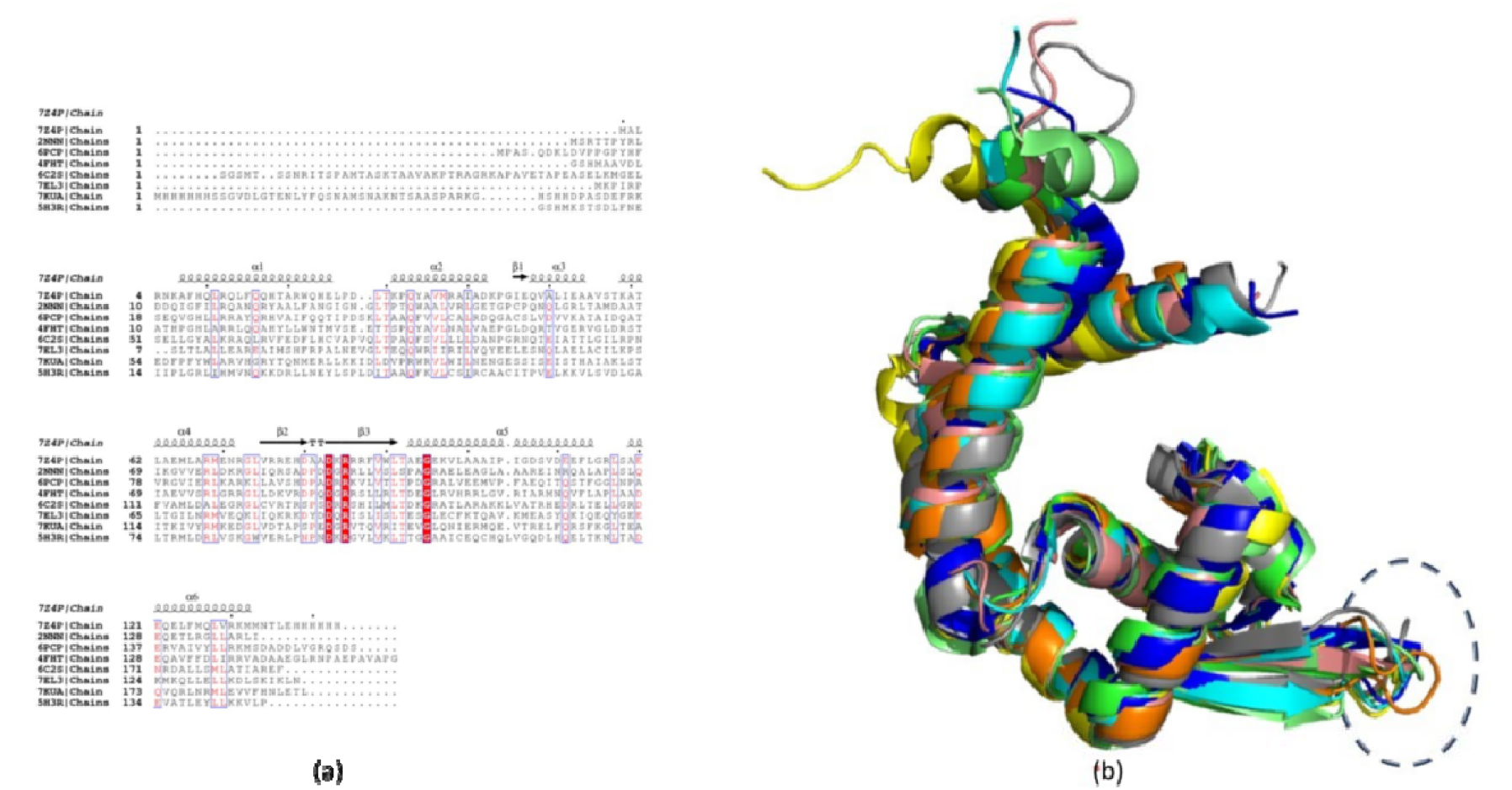
(a) Multiple sequence alignment and secondary structure assignment: Secondary structure elements of HosA protein (8PQ4) and structural homologues are displayed above the sequences: *α*-helices as medium squiggles and *β*-strands as arrows. Amino acids that appear in white characters in a red background are identical in all aligned proteins, while those in red characters and in blue frames are conserved in the majority of proteins. (b) Structural superposition of HosA (orange) with similar structures 2NNN (green), 4FHT (cyan), 6C2S (yellow), 6PCP (salmon),7EL3 (blue), 7KUA (grey) and 5H3R (magenta). The proteins are shown by cartoon representation and dashed line encircles the loop region.

### 3.3 Adaptive Poisson-Boltzmann Solver (APBS) analysis

APBS analysis was carried out at neutral pH in PDB2PQR/APBS server to visualize the solvent accessible surface area (SASA) charges. The monomeric protein showed contrasting opposite charges on either side (Supplementary Fig. S1a). This charge based axial dichotomy can be the driving force behind crystallographic dimer formation as predicted from PISA analysis. Interestingly, the base surface of the negatively charged side has accumulated positively charged residues (Supplementary Fig. 1a). While the positively charged side has a bottom of mostly positively charged and some neutral amino acids (Supplementary Fig. S1a). Therefore, the bottom base of the dimeric complex form a wide area with dense positive charges (Supplementary Fig. S1b), ideal to interact with negatively charged DNA. These findings provide valuable insights into the surface properties of the protein, which can aid in understanding its function and interactions with other molecules along with DNA.

## Acknowledgements

Authors thank Rukmini R and Malesh K for technical support during data collection in X-ray facility, CSIR-CCMB, Hyderabad, India. Also, Dr. Kavyashree Manjunath, InStem, Bangalore, India is duly acknowledged for her suggestions during data processing. Dr. Manjunatha NK, SVCE, Bangalore, India is thanked on communications for diffraction. We also thank IOE, University of Mysore, Mysuru, India for support for crystallization accessories. MKS thank O.E. och Edla Johanssons vetenskapliga stiftelse.

